# Cortical changes associated with parenthood are present in late life

**DOI:** 10.1101/589283

**Authors:** Edwina R Orchard, Phillip GD Ward, Francesco Sforazzini, Elsdon Storey, Gary F Egan, Sharna D Jamadar

## Abstract

Early parenthood results in changes in cortical thickness in regions related to parental care. However, the enduring effects of this period on the structure of the human brain, and cognition in late-life, is unknown. In an elderly sample, we examined the relationship between the number of children parented (here, 1-6 children) and cortical thickness in 267 males (74.0 ±3.5 years) and 231 females (73.8±3.5 years). We also compared cognition and cortical thickness between parents of one child and non-parents, in n=36 males (73.4±3.7 years), and n=46 females (72.8±3.3 years). We obtained a positive relationship between number of children parented and verbal memory performance, showing increasing memory performance with number of children. For mothers, number of children positively correlated with cortical thickness in the parahippocampal gyrus and negatively correlated with regions of the visual cortex. Mothers of one child showed thinner cortical thickness in the dorsolateral prefrontal cortex and visual cortex compared with childless women. Fathers of one child showed thinner cortical thickness in the anterior cingulate cortex and thicker cortical thickness in the temporal pole compared with childless men. Our results are the first to reveal distributed differences in cortical thickness related to parenthood that are evident beyond the postpartum period. Our findings overlap substantially with the areas found to be altered across pregnancy and the postpartum period, suggesting that neural changes associated with early parenthood persist into older age, and are potentially cognitively beneficial.

## Introduction

New parents face an endless series of novel challenges to ensure the survival of their offspring. In addition to their own personal needs and existing responsibilities, new parents must confront persistent demands for care and protection of their infant. As children grow, new challenges arise: swapping nappies and baby bottles for school lunches and soccer practice. The increased environmental complexity of parenthood continues for years - even decades, and is increased by each subsequent child; requiring simultaneous care across different stages of dependency. Successfully navigating these challenges requires rapid behavioural change and skill acquisition, driven by a combination of neuroendocrine and experiential factors (Stolzenberg & Champagne, 2016). The lasting impact of parenthood on the human brain is unknown, with research in this field still in its infancy.

The overwhelming majority of studies investigating the parental brain are preclinical, and almost all have been conducted in females. In rodents, hormonal changes during pregnancy prime the mother for the onset of novel, pup-directed maternal behaviours, which are triggered after parturition in the presence of pups (Rosenblatt, 1994). Converging evidence in non-human animals suggest that neural adaptations support maternal behaviours, including changes to dendritic morphology, cell proliferation and gene expression (Hillerer, Neumann, Couillard-Despres, Aigner, & Slattery, 2014; Kinsley & Amory-Meyer, 2011). In the rodent, neural changes centre around the amygdala, parietal cortex, prefrontal cortex, and the oxytocin-producing medial preoptic area (MPOA) of the hypothalamus (Fleming & Korsmit, 1996). Changes to the MPOA are crucial for the development of maternal behaviour, and interruption to the function of this region results in complete absence or cessation of maternal behaviours (Numan, Corodimas, Numan, Factor, & Piers, 1988; Numan & Insel, 2003; Numan & Numan, 1991). Taken together, these studies suggest that the mammalian brain undergoes substantial restructuring to accommodate the demands of parenthood.

Although well established in rodents, the impact of reproductive experience on the human brain is less clear. Studies on human parenthood are largely infant-centric, rather than parent-centric. In other words, most existing studies focus on the parents’ response to the infant, rather than more generalised changes to the parental brain. Most studies use functional magnetic resonance imaging (fMRI) to examine parents’ brain response to infant pictures, movies, or sounds of infant cries (see Swain, Lorberbaum, Kose, & Strathearn, 2007 for review). Several brain areas in human adults consistently activate in response to infant cues, which has been labelled the ‘parental caregiving network’ (Feldman, 2015). The parental caregiving network encompasses a number of interconnected sub-networks that together define the social brain, including the motivation-reward, empathy, emotional regulation and mentalising networks (Feldman, 2015). The parental care-giving network is differentially activated in mothers and fathers in response to infant pictures, movies and cries, depending on the sex of the individual, and whether they are a primary or secondary caregiver (Abraham et al., 2014). Neural changes associated with parenthood can be enhanced through greater interaction with the child (i.e., increased time spent in childcare), suggesting that this network can be flexibly activated depending on the environmental context (Abraham et al., 2014; Feldman, 2015).

A surprisingly small number of studies (five) have examined structural changes associated with parenthood in humans, and most (four of five) have focused on changes in females but not males. These studies are small (sample sizes between 8-25), which may explain the inconsistent patterns of results. In mothers, Oatridge et al. (2002) found that during pregnancy, total brain size decreased and ventricular volume increased compared with pre-conception, and returned to pre-conception size by 24 weeks post-partum. Kim et al. (2010) found that new mothers showed increased grey matter volumes throughout the cerebral cortex four months post-partum, compared with 1-month post-partum. Increased grey matter volume in the midbrain at four months post-partum predicted the mother’s positive opinions of her own baby. In a follow-up study, Kim, Dufford, & Tribble (2018) found increasing grey matter thickness in mothers across the first 6 months post-partum, and identified a positive correlation between cortical thickness and parental self-efficacy. Conversely, Hoekzema et al. (2017) showed that pregnancy resulted in substantial *decreases* in grey matter thickness, particularly in networks subserving social cognition and theory of mind. Reduced grey matter in mothers across pregnancy predicted increased mother-infant attachment and was sustained for at least two years post-partum. Fatherhood also modulates plasticity in regions related to parental care. Kim et al. (2014) found regional grey matter volume changes (both increases and decreases) in fathers at four months post-partum compared with one month post-partum. Together, the results of these studies show an inconsistent pattern of increases and decreases in cortical thickness in early parenthood, which may suggest that early parenthood involves dynamic and stage-specific neural reorganisation, and contains periods of alternating cortical thinning and thickening. Consequently, the neurobiological effects of parenthood on the human brain may be subtle and require larger studies to characterise accurately.

It is not yet clear how long parenthood-related changes to the brain are sustained after the very early parental period (parturition), but preliminary evidence from preclinical models suggest that the changes may be permanent. Strikingly, preclinical evidence suggests that motherhood may convey a protective effect against dementia. Gatewood et al. (2005) found reduced hippocampal amyloid deposits and attenuated memory decline in older mother rats, compared with older virgin rats. Thus, reproductive experience appears to benefit rodents into senescence, with hormonal and environmental changes that interact to produce a parental brain that is healthier, more flexible and resistant to age-related decline (Gatewood et al., 2005; Kinsley et al., 1999; Macbeth & Luine, 2010). Determining whether these neuroprotective effects are evident in humans, as well as rodents and other mammals, would contribute to our understanding of the protective advantages of parenthood on neuroanatomical changes and preserved cognition in late-life.

No study has investigated the enduring effects of parenthood on the structure and function of the human brain beyond two-years post-partum. Here, we report the ‘dose’ effects of parenthood by examining the relationship between cortical thickness and the number of children a person has parented, in a large healthy aged sample that forms part of the ASpirin in Reducing Events in the Elderly (ASPREE) clinical trial; specifically, the ASPREE-NEURO sub-study (Ward et al., 2017). We also report long-term differences in brain structure between parents and non-parents that are sustained across the lifespan and into older age. We hypothesised that regional cortical thickness differences would differ between parents and non-parents and be associated with the number of children. For both sexes, early parenthood is likely to involve environmental, cognitive and psychosocial changes, and therefore we anticipated that changes found in the maternal brain would also be evident in the paternal brain albeit to a lesser extent (Storey, Walsh, Quinton, & Wynne-Edwards, 2000).

## Methods

### Participants

All participants were enrolled in ASPREE-NEURO, which is a sub-study of the ASPirin in Reducing Events in the Elderly (ASPREE) clinical trial. (Baseline characteristics of the full ASPREE sample are reported in McNeil et al., 2017). Imaging data was acquired as part of the ASPREE-NEURO sub-study (Ward et al., 2017), which aimed to determine the effect of low-dose aspirin on a range of MRI biomarkers (cerebral microbleeds, white matter hyperintensities) over a three-year period, in neurotypical adults aged 70-yrs and over. We included data from baseline (prior to study medication), and scans taken one year and three years later. Older adults in Australia were eligible to participate if aged 70-years and over, had no history of occlusive vascular disease, atrial fibrillation, cognitive impairment or disability; were not taking antithrombotic therapy, and did not have anaemia or a diagnosis likely to cause death within five years. At study entry, each participant had a Modified Mini Mental Status Examination (3MS; Teng & Chui, 1987) score of at least 78/100.

Our participants (N=573) were aged 70-88 years. Data from 21 participants contained incomplete MRI scans and were discarded. Data from five participants were excluded due to the presence of neurological conditions (history of brain cancer (n=1), Parkinson’s disease (n=4)). In the final sample, data from 547 subjects aged 70-88 years (mean age [+/-stdev] = 73.9+/-3.5years) were retained for the first timepoint, including n=287 males (74.0 ±3.6 years), and n=260 females (73.7 ±3.4 years). Participant attrition across the three timepoints is given in Supplementary Table 1.

As part of a health outcomes questionnaire, ASPREE-NEURO participants were asked “How many children do you have?”. More detailed parenthood data (e.g. biological/adopted children, grandparenthood, primary/secondary caregiver, absence from children etc.) were not collected. The number of children parented by males and females in our sample is shown in Supplementary Figure 1. Participants with seven or more children (n=6) were excluded from analyses due to their low frequency.

### Data Acquisition

#### Cognitive Tasks

At baseline, participants completed a 30-minute battery of cognitive tasks, including the Controlled Oral Word Association Test (COWAT; Ross, 2003), Hopkins Verbal Learning Test-Revised (HVLT-R; Brandt & Benedict, 2001), Symbol Digit Modalities Test (SDMT; Smith, 1982), Color Trails (D’Elia, Satz, Uchiyama, & White, 1996), Stroop Test (Victoria version; Troyer, Leach, & Strauss, 2006) and Center for Epidemiological Studies Depression Inventory (CES-D; Radloff, 1977).

#### Image Acquisition and Preparation

All MRI scans were obtained on a 3T Siemens Skyra MR scanner (Erlangen, Germany) at Monash Biomedical Imaging, Melbourne, Australia. ASPREE-NEURO sequence protocols are described in detail elsewhere (Ward et al., 2017). Here, we used T1 and T2 data. T1-weighted magnetization-prepared rapid gradient-echo (MPRAGE) images were acquired (TR=2300 ms, TE=2.13 ms, TI=900 ms, matrix size=256×240×192, bandwidth=230 Hz/pixel). T2-Fluid attenuated inversion recovery (FLAIR) images (TR=5000ms, TE=395ms, TI=1800ms, matrix size=256×240×160, bandwidth=781Hz/pixel) were also acquired, to highlight gross white matter abnormalities and detect white matter hyper-intensities.

To extract cortical thickness values, MR images were segmented using the ‘recon-all’ function of FreeSurfer v5.3.0 (Dale, Fischl, & Sereno, 1999). To optimise the extraction of the pial surface (cerebrospinal fluid-grey matter boundary), both T1 MPRAGE and T2 FLAIR scans were used. Cortical thickness values were extracted for 34 regions (plus mean thickness) for each hemisphere using the Desikan-Killiany atlas, producing 70 cortical thickness values in total (2 hemispheres × 34 regions + 2 × hemisphere means; Desikan et al., 2006).

### Data Analysis

A step-wise regression was conducted to control for the effects of age, education, systolic and diastolic blood pressure, BMI, and cholesterol. Z-scores of the residuals were taken and used throughout the subsequent analyses. Regressors were selected based on previous studies reporting correlations between these factors and cortical thickness (Seo et al., 2011; Veit et al., 2014). Although cortical thickness measurements were computed at three timepoints, the effect of parenthood in late life was not expected to change across the three year measurement period. Rather, any differences in the relationship between cortical thickness and parenthood across time was assumed to be attributable to measurement error. Therefore, to control for false positives due to measurement error, the mean cortical thickness for each participant was calculated from the three measurements for each brain region.

The ‘dose’ effects of the number of children on cortical thickness were tested using Spearman’s correlation. The analysis was performed on all 70 thickness measures for 498 elderly subjects, n=267 male (74.0 ±3.5 years) and 231 female (73.8±3.5 years) participants who were parents of between one and six children. To test the hypothesis that cortical thickness would differ between parents and non-parents, for both sexes, the cortical thickness of 70 brain regions were compared between individuals with one child and individuals with no children in 82 elderly subjects, including males (n=36; 73.4 ±3.7 years) and females (n=46; 72.8 ±3.3 years). The effects of parenthood were quantified using Cohen’s *d*. Significance was estimated using a permutation test (10,000 permutations of group assignments).

Data analysis for the cognitive tasks mirrored the analysis structure of the MRI data. All cognitive tasks (COWAT, HVLT-R, SDMT, Color Trails, Stroop and CES-D) were compared between participants who were childless (non-parents) and those who were parents to one child. The relationship between all cognitive tasks and number of children (range 1-6) was also examined. As this study is highly novel and exploratory in nature, no correction for multiple comparison was applied to avoid type II errors (Armstrong, 2014).

## Results

For mothers, three regions showed a significant relationship with number of children (1-6 children; Table 1; Figure 1). The left pericalcarine sulcus and cuneus showed a negative correlation with number of children, with cortical thickness decreased as number of children increased. The right parahippocampal gyrus showed a positive correlation with number of children (Figure 2). For fathers, the thickness of the right fusiform gyrus and the left temporal pole showed a positive correlation with number of children (1-6 children; Figure 3).

**Table 1:**
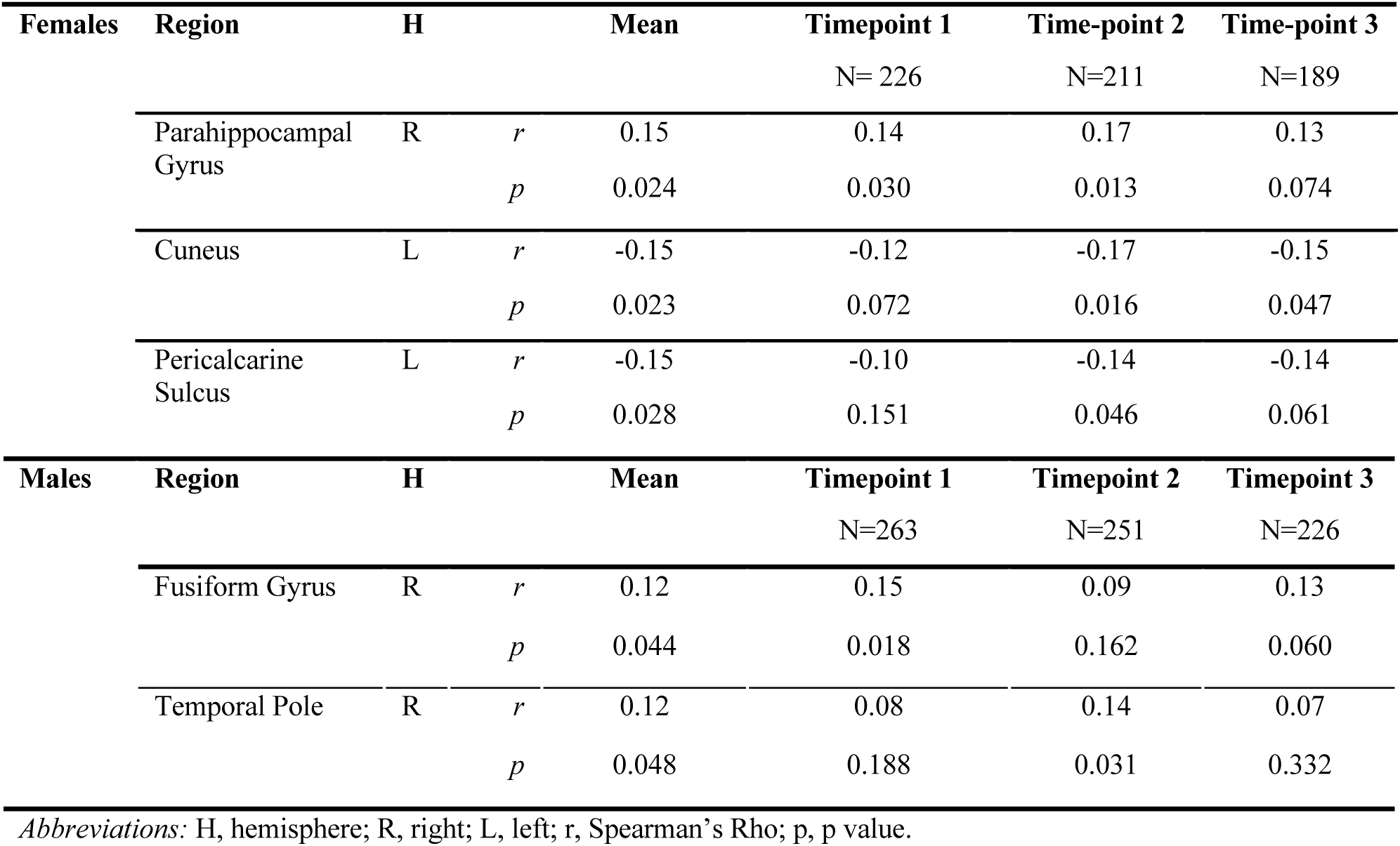
Relationship between number of children and cortical thickness for females and males.

**Figure 1:**
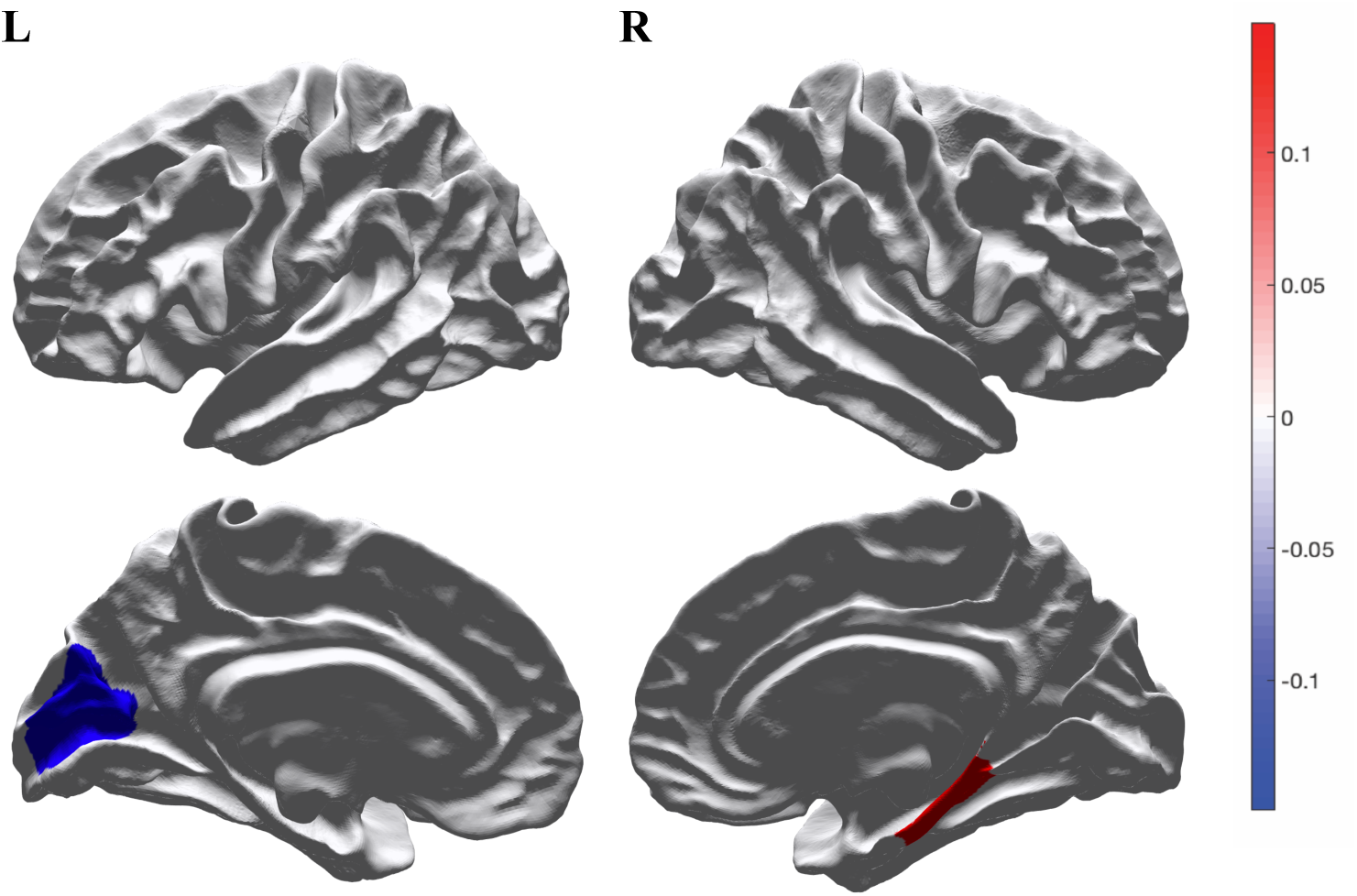
Spearman’s Rho for brain regions showing a significant relationship between cortical thickness and number of children for female. Regions highlighted in red (right parahippocampal gyrus) depict a positive relationship, regions highlighted in blue (left pericalcarine sulcus and cuneus) depict a negative relationship.

**Figure 2:**
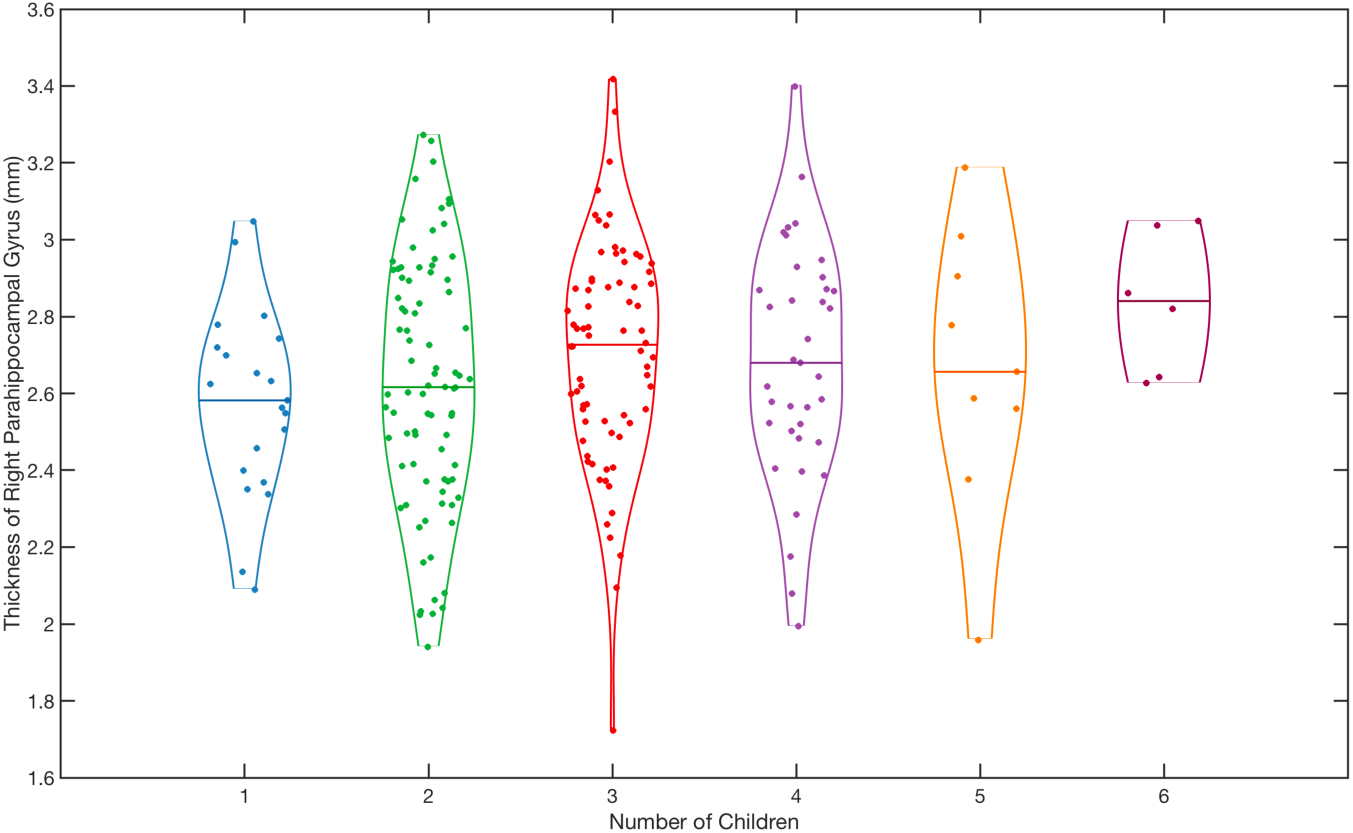
Cortical thickness of the right parahippocampus (mm) for participants with one to six children, showing a positive relationship between number of children and thickness, Spearman’s Rho = 0.15, p=0.024

**Figure 3:**
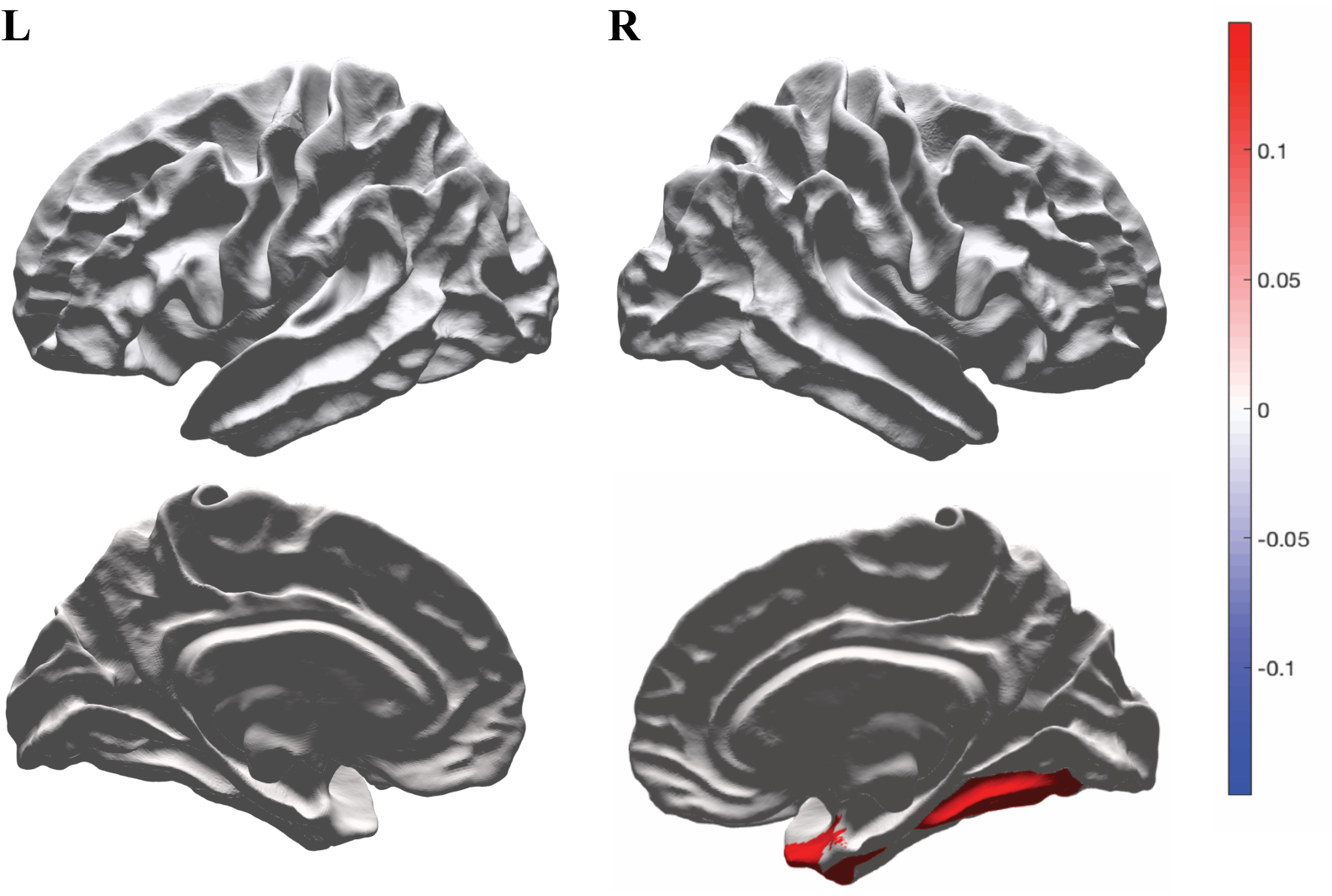
Spearman’s Rho for brain regions showing a significant relationship between cortical thickness and number of children for males. Regions highlighted in red (right fusiform gyrus and temporal pole) depict a positive relationship.

In females, two regions showed consistent differences in cortical thickness between mothers of one child and childless women (non-mothers). The left dorsolateral prefrontal cortex and the right pericalcarine sulcus showed thinner grey matter in mothers compared with non-mothers (Table 2; Figure 4). In males, two regions showed significant differences in cortical thickness between fathers of one child and those who had not fathered children. The left anterior cingulate showed thinner grey matter in fathers compared with non-fathers, and the right temporal pole showed thicker grey matter in fathers compared with non-fathers (Figure 5).

**Table 2:**
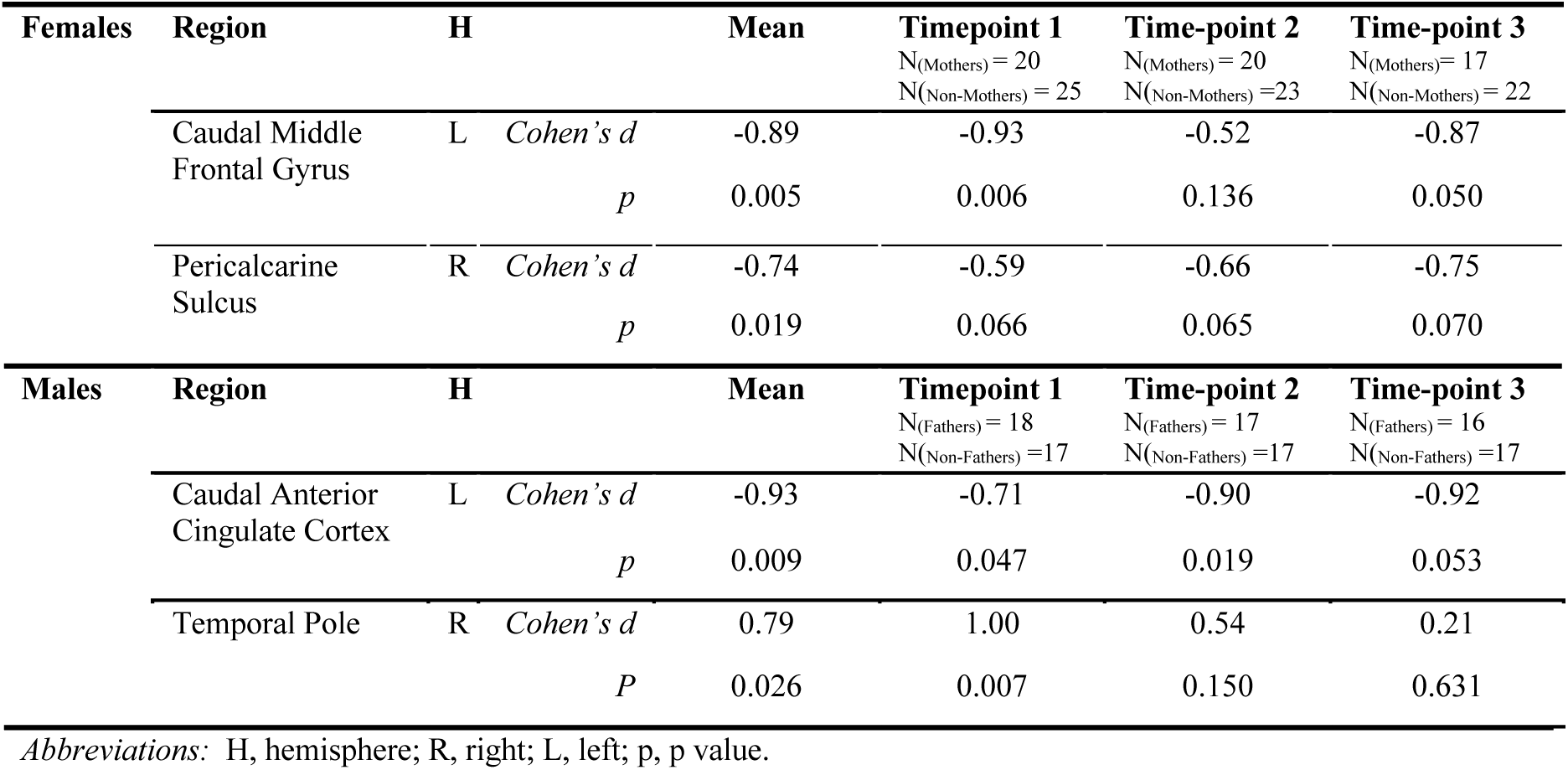
Cortical thickness differences between parents (one child) and non-parents for males and females.

**Figure 4:**
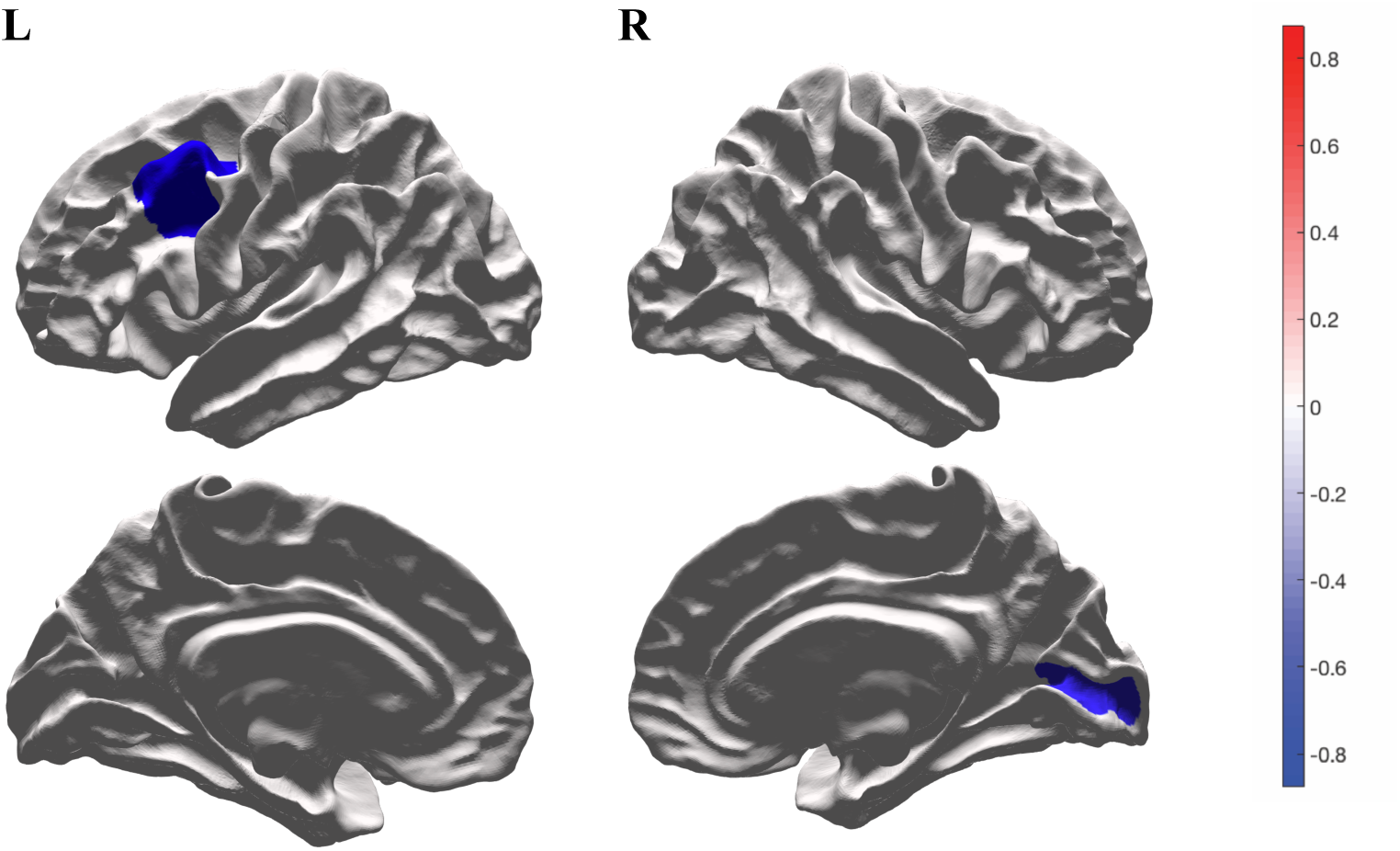
Cohen’s d for the difference between females with one child and females with zero children. Regions highlighted in blue (left caudal middle frontal/DLPFC, right pericalcarine sulcus) depict a negative Cohen’s d and thinner GM in mothers, compared with non-mothers.

**Figure 5:**
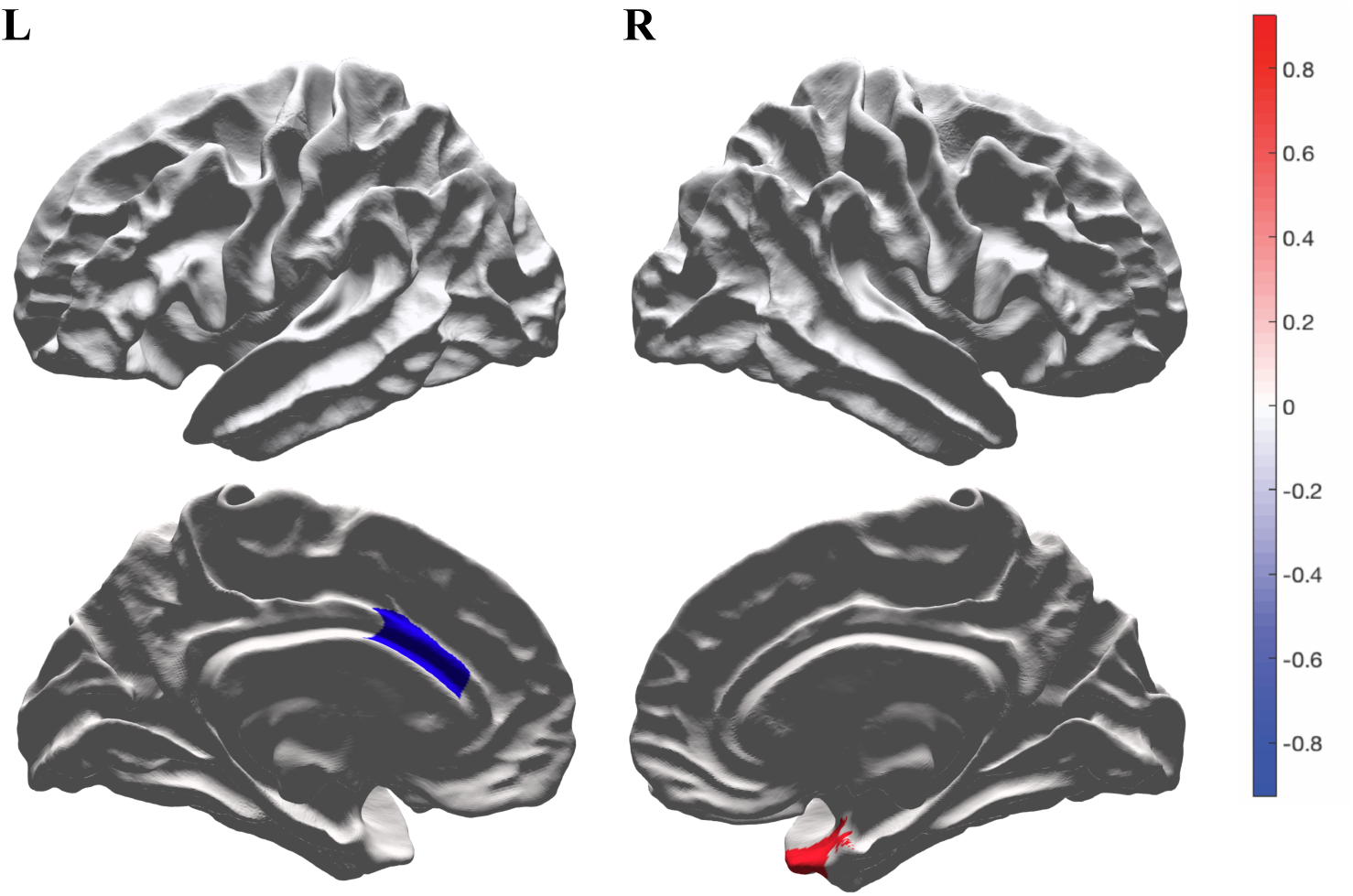
Cohen’s d for the difference between males with one child, to males with zero children. Regions highlighted in red (right temporal pole) depict a positive Cohen’s d and thicker GM in fathers, regions highlighted in blue (left caudal anterior cingulate cortex) depict a negative Cohen’s d and thinner GM in fathers compared with non-fathers.

No significant differences were obtained between childless and parental participants of either sex for any of the cognitive tests. In males, no relationship was found between any cognitive test and parenthood. Females showed a marginally significant positive relationship between parity and delayed recall scores on the Hopkins Verbal Learning Test - Revised (HVLT-R), *r*=0.17, *p*=0.011, such that women with more children showed better HVLT-R memory outcomes.

## Discussion

In this study, we investigated differences in cortical thickness in parenthood that are long-term and persist into older age. No study has ever examined the effect of parenthood on brain structure in humans beyond two years post-partum. The anatomical distribution of the observed effects has previously been confined to human pregnancy and early parenthood in structural (Hoekzema et al., 2017; Kim et al., 2018; Kim et al., 2010; Kim et al., 2014) and functional studies of parenthood (see Swain et al., 2007 for review).

We found a positive relationship between number of children and cortical thickness in the parahippocampal gyrus of elderly mothers. This result is consistent with findings in early parenthood, where grey matter changes are seen in the parahippocampal gyrus during pregnancy (Hoekzema et al., 2017) and the postpartum period (Kim et al., 2010). The parahippocampal gyrus is associated with memory, which is compatible with our cognitive results, where memory outcomes were improved in elderly women with more children. Motherhood appears to be associated with a small but reliable improvement in memory outcomes for women, which is sustained into older age. This result is consistent with evidence from early parenthood, where memory is impaired during pregnancy (Davies, Lum, Skouteris, Byrne, & Hayden, 2018), returns to pre-pregnancy levels in the postpartum period (Buckwalter, Buckwalter, Bluestein, & Stanczyk, 2001; Macbeth & Luine, 2010), and is enhanced in late parenthood (Gatewood et al., 2005).

Our result is consistent with results seen in preclinical models. Spatial learning and memory are improved in rodents with reproductive experience, and this effect is stronger in multiparous mothers (mothers with more than one pregnancy; Kinsley & Lambert, 2008; Macbeth & Luine, 2010). This cognitive benefit persists into late-life for the parous rats, who have increased hippocampal long-term potentiation, enhanced memory capacities and fewer signs of brain aging compared with aged nulliparous females; an effect that is enhanced with multiparity (Gatewood et al., 2005). Our findings, that both memory performance and parahippocampal thickness increase with multiparity in humans, are consistent with these reports and suggest that parenthood positively impacts human cognition and may enhance neuroplasticity in late-life.

Compared with childless women, those with one child showed thinner grey matter in the left dorsolateral prefrontal cortex. The dorsolateral prefrontal cortex is involved in high-order cognitive functions including executive functioning, cognitive control and emotional regulation (Alvarez & Emory, 2006; Wager, Davidson, Hughes, Lindquist, & Ochsner, 2008). Neuroimaging data highlights the importance of the prefrontal cortex in parenting behaviours, as this region is activated in almost every fMRI study of human mothers’ responses to infant stimuli (Swain et al., 2007). Increased brain activity in a mother’s prefrontal cortex in response to her own baby (compared to an unfamiliar baby) is positively associated with maternal sensitivity and higher quality of mother-infant interactions (Atzil, Hendler, & Feldman, 2011; Musser, Kaiser-Laurent, & Ablow, 2012). Conversely, decreased fMRI activation in the prefrontal cortex is correlated with difficulties adjusting to parenthood (Laurent & Ablow, 2011; Swain et al., 2008). Neural changes in early motherhood are likely to have evolved to assist in the dual survival of parent and young (Kinsley, Franssen, & Meyer, 2011). Three recent studies have shown maternal cortical thickness changes in the prefrontal cortex across pregnancy and the early post-partum period (Hoekzema et al., 2017; Kim et al., 2018; Kim et al., 2010). In all three of these studies, structural change was associated with increased maternal sensitivity (attachment; parental self-efficacy; positive opinions about their baby). Our results indicate that parenthood-related changes in the cortical thickness of the dorsolateral prefrontal cortex are sustained across the lifespan.

Our results show thinner cortical thickness of the visual cortex (pericalcarine and cuneus) for females. These results appear in both hemispheres across both analyses (childless versus one child, and number of children). Interactions with children provide a rich array of sensory experiences for mothers. Interactions involving multi-sensory cues (e.g., visual, auditory, tactile) between mothers and infants throughout the postpartum period, and continuing into late-parenthood, may alter the structure and function of the maternal sensory regions over time (Kim et al., 2018). In rodents, neural reorganisation of the olfactory bulb, parietal lobe, and somatosensory cortex were associated with the expression of maternal behaviours during the postpartum period (Kinsley & Lambert, 2008). Our findings of cortical thickness changes in regions involved in visual information perception and processing during the early postpartum period are consistent with earlier findings in human mothers, (grey matter volume; Kim et al., 2018) and enhanced brain activation in response to own infant cues (Nishitani, Kuwamoto, Takahira, Miyamura, & Shinohara, 2014).

Compared with childless men, fathers showed thinner grey matter in the left caudal anterior cingulate cortex and thicker grey matter in the right temporal pole. The caudal anterior cingulate and temporal pole are both involved in social cognition, including emotional regulation, empathy and theory of mind (Etkin, Egner, & Kalisch, 2011; Olson, Plotzker, & Ezzyat, 2007). Social cognition is acknowledged as a hallmark of the human parental brain, especially relevant for pre-verbal infants (Lenzi et al., 2009). Theory of mind abilities, including empathy, contribute to successful social interactions and are important for human survival. Our results support previous studies of early parenthood that found grey matter volume changes in the anterior cingulate cortex associated with early parenthood in mothers (Hoekzema et al., 2017; Kim et al., 2010) and fathers (Kim et al., 2014), and suggests they persist into late-life. Fathers also showed a positive correlation between number of children and cortical thickness for the fusiform gyrus and temporal pole. These correlations are close to the threshold of significance and require further studies to comprehensively characterise this effect.

We argue that the structural brain changes reported in this study, which endure throughout the lifespan, are related to the environmental changes of parenthood. While the drastic hormonal shifts of pregnancy may play an initial role in mothers, the hormonal milieu of pregnancy returns to a pre-pregnancy state following parturition, at which time many hormonal and biological changes are resolved, i.e., uterus size, lactation, etc (Henry, 2016). In our elderly sample the hormonal exposure of pregnancy occurred three or more decades earlier. Conversely, the environmental complexity of parenthood endures throughout the entire parenting experience, possibly throughout the life-span of a parent, and compounds with increasing number of children. The neural impact of parenthood is evident in non-human mammals from environmental changes alone. Exposure to unfamiliar pups for ten minutes for four days was sufficient to induce neural changes and improve spatial memory for non-parental male rats (Everette et al., 2007; Everette et al., 2006). This suggests that the mere presence of offspring produces neural change in the absence of pregnancy-induced hormonal changes.

Child-rearing provides parents of both sexes with rich sensory stimulation and increased environmental complexity that may be comparable to traditional concepts of an enriched physical environment. It has long been recognised that complex, or ‘enriched’ environments lead to improved outcomes and neuroprotective effects against ageing. Enriched environments stimulate synaptogenesis (Diamond, Krech, & Rosenzweig, 1964) and more complex dendritic branching (Volkmar & Greenough, 1972), and may provide a protective effect against brain ageing (Speisman et al., 2013). For instance, male and female rats housed in enriched conditions develop greater cortical depth (*ex vivo*) compared with littermates in impoverished settings. Furthermore, cortical depth reported in post-partum rats housed in environmentally impoverished conditions resembled that of virgin rats housed in enriched environments, suggesting the maternal brain was stimulated in a manner similar to typical enrichment protocols (Diamond, Johnson, & Ingham, 1971). Marmoset fathers involved in the rearing of offspring also experience similar brain changes to non-parent animals living in enriched environments, and this effect is mediated by the amount of father-infant contact (Kozorovitskiy, Hughes, Lee, & Gould, 2006). This reflects literature on the subject in humans and suggests that the amount of parent-infant contact is associated with the degree of neuroanatomical change (Abraham et al., 2014).

In the present study, cortical thickness differences between parents and non-parents were evident in both sexes. This novel finding implicates the parental environment, which exists for both sexes. However, the neuroanatomical changes associated with parenting multiple children were far more significant for mothers, with only marginal effects identified in fathers. This may reflect differences in care-giving responsibilities between the sexes: the “silent generation” cohort examined in this study most likely included participants who were predominantly in ‘traditional’ care-giving arrangements, with roles and responsibilities as primary care-giving mothers and ‘bread-winning’ fathers (Broomhill & Sharp, 2004). The social structure of the silent generation may have limited the cumulative impact of parenthood for the males in this cohort. The current findings are the first to show a relationship between number of children and grey matter thickness in the ageing human brain. Future studies to investigate neurobiological changes across parenting of multiple children in humans are required, as there are currently no such studies in humans at any life-stage (Maupin, Roginiel, Rutherford, & Mayes, 2016).

The diverse and subtle changes in cortical grey matter thickness measured in this study point to a rich and multi-layered process of cortical plasticity that begins during pregnancy (in females) and early parenthood (males and females), and is sustained across the lifespan. The physiological processes that can alter cortical thickness include changes in the number of synapses, glial cells or neurons, morphological changes to dendritic structure and axonal myelination, and vasculature changes - including angiogenesis - that influence blood volume and perfusion. While the study design precludes identification of the neurobiological mechanisms that underpin the observations, the novelty and social importance of the findings highlight the need for further studies: *in vivo*, to examine the microstructural, functional and neurovascular characteristics of parenthood; and post-mortem stereological, to determine the cellular and morphological origins of the cortical thickness changes.

The data used in this study were acquired as part of a larger study with broader aims. Detailed demographic information related to parenthood was not obtained from the participants, and so information about the parenting experiences is not available. For example, we do not know about birth details, whether children were adopted or survived beyond birth, the parental style, and whether participants are involved in the care of grandchildren. Many of these unknown factors, such as birth and feeding methods, hormone levels and parental sensitivity, potentially impact maternal neuroplasticity(Kim, Strathearn, & Swain, 2016). Future studies should control for these factors, as well as examine the potential effects of grandparenthood and other opportunities for interaction with children during later life.

## Conclusion

This study is the first examination of the effect of parenthood on the ageing human brain. The results are consistent with published neuroanatomical and behavioural findings in early parenthood, and reveal distributed differences in cortical thickness related to parenthood that are evident beyond the post-partum period. The findings suggest that grey matter changes associated with early parenthood, for both sexes, endure throughout the life-span and into older age. Our understanding of the structural changes related to human parenthood remains in its infancy, even during pregnancy and the early post-partum period. The neuroscience of parenthood is a fertile area for further research to identify the critical time periods of parenthood when the brain is highly plastic, and examine longitudinal changes beyond the immediate post-partum months. The nascent field of the neuroscience of parenting has the potential to elucidate the neurobiological and neuroanatomical impact of the human reproductive experience, which may provide unique insights into mechanisms of neuroplasticity.

## Acknowledgements

This work was supported by an Australian National Health and Medical Research Council Project Grant (APP1086188). ASPREE was supported by the National Institutes of Health (grant number U01AG029824); the National Health and Medical Research Council of Australia (grant numbers 334047, 1127060); Monash University (Australia) and the Victorian Cancer Agency (Australia). WO, PW, GE and SJ are supported by the Australian Research Council (ARC) Centre of Excellence for Integrative Brain Function (CE140100007). SJ is supported by an ARC Discovery Early Career Researcher Award (DE150100406). The Principal ASPREE study is registered with the International Standardized Randomized Controlled Trials Register, ASPirin in Reducing Events in the Elderly, Number: ISRCTN83772183 and clinicaltrials.gov number NCT01038583. ASPREE-Neuro trial is registered with Australian New Zealand Clinical Trials Registry ACTRN12613001313729.

## Supplementary

**Supplementary Table 1:**
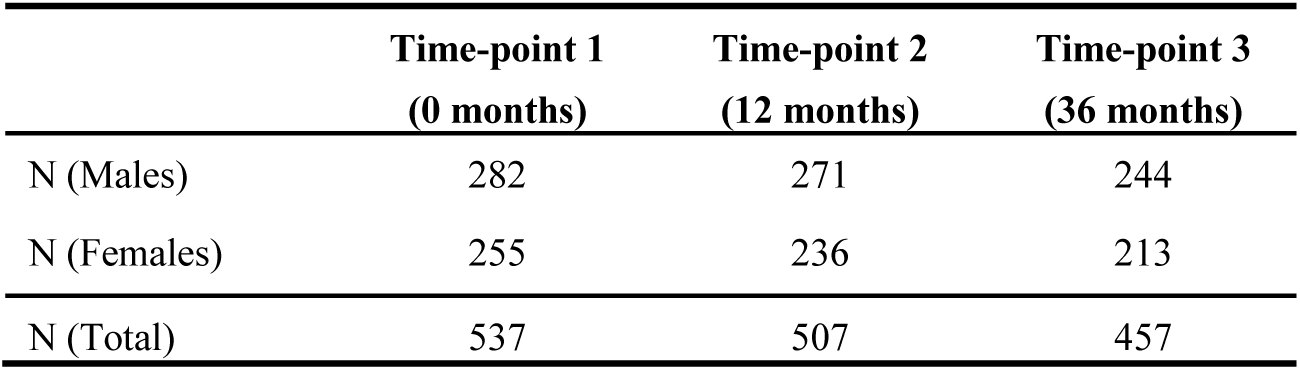
Participant numbers in the study cohort at 0, 12 and 36 months showing a 15% attrition across three years.

|  | Time-point 1<br>(0 months) | Time-point 2<br>(12 months) | Time-point 3<br>(36 months) |
| --- | --- | --- | --- |
| N (Males) | 282 | 271 | 244 |
| N (Females) | 255 | 236 | 213 |
| N (Total) | 537 | 507 | 457 |

**Supplementary Figure 1:**
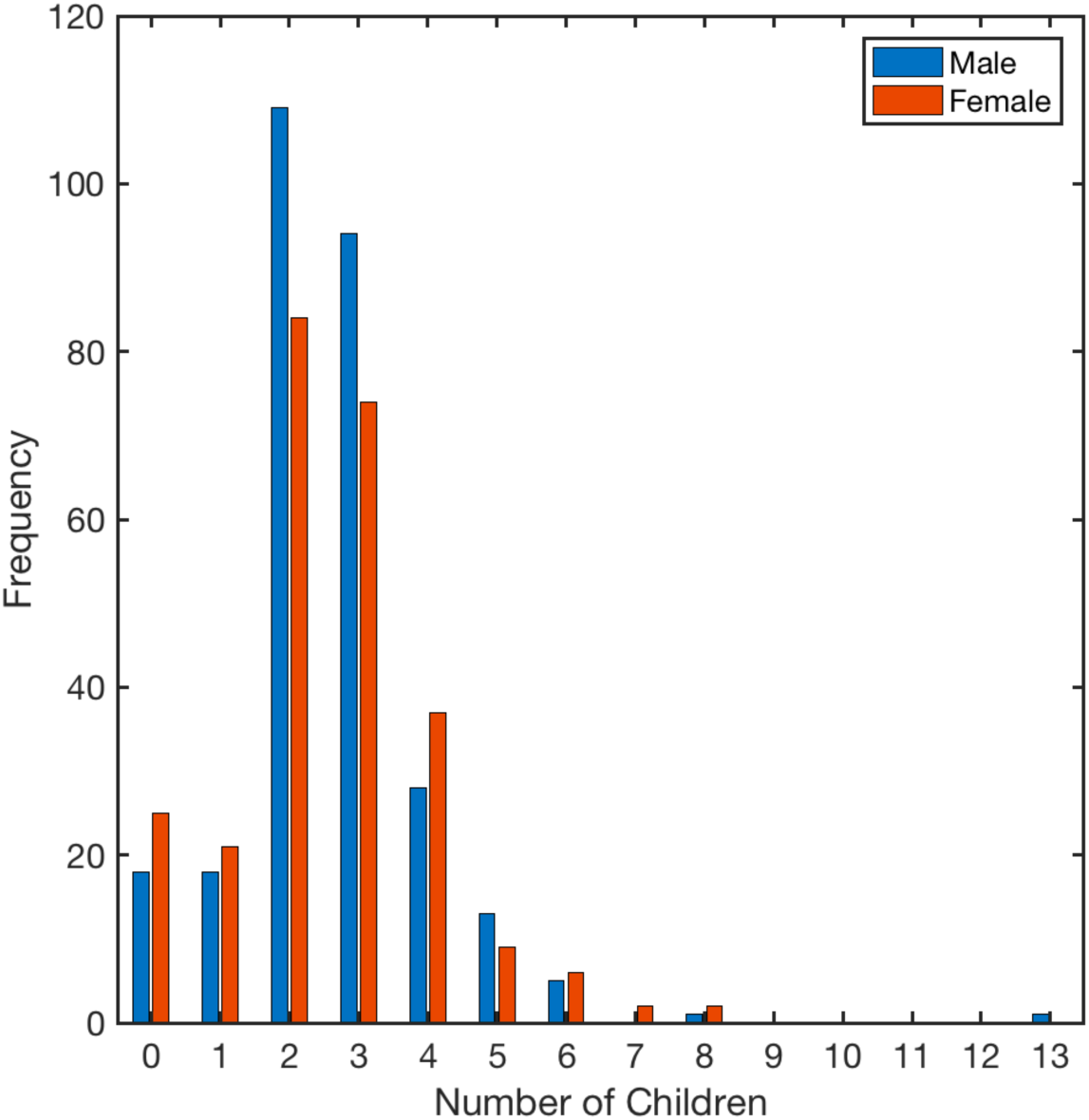
Number of children per participant, split by gender

## References

Abraham, E., Hendler, T., Shapira-Lichter, I., Kanat-Maymon, Y., Zagoory-Sharon, O., & Feldman, R. (2014). Father’s brain is sensitive to childcare experiences. Proceedings of the National Academy of Sciences, 111(27), 9792–9797. doi:10.1073/pnas.1402569111

Alvarez, J. A., & Emory, E. (2006). Executive function and the frontal lobes: a meta-analytic review. Neuropsychology review, 16(1), 17–42.

Armstrong, R. A. (2014). When to use the Bonferroni correction. Ophthalmic and Physiological Optics, 34(5), 502–508.

Atzil, S., Hendler, T., & Feldman, R. (2011). Specifying the neurobiological basis of human attachment: brain, hormones, and behavior in synchronous and intrusive mothers. Neuropsychopharmacology, 36(13), 2603.

Brandt, J., & Benedict, R. H. (2001). Hopkins verbal learning test-revised: professional manual: Psychological Assessment Resources.

Broomhill, R., & Sharp, R. (2004). The changing male breadwinner model in Australia: A new gender order? Labour & Industry: a journal of the social and economic relations of work, 15(2), 1–23.

Buckwalter, J. G., Buckwalter, D. K., Bluestein, B. W., & Stanczyk, F. Z. (2001). Pregnancy and postpartum: changes in cognition and mood. In Progress in brain research (Vol. 133, pp. 303–319): Elsevier.

D’Elia, L., Satz, P., Uchiyama, C., & White, T. (1996). Color Trails Test. Odessa, FL: Psychological Assessment Resources. Inc Google Scholar.

Dale, A. M., Fischl, B., & Sereno, M. I. (1999). Cortical surface-based analysis: I. Segmentation and surface reconstruction. Neuroimage, 9(2), 179–194.

Davies, S. J., Lum, J. A., Skouteris, H., Byrne, L. K., & Hayden, M. J. (2018). Cognitive impairment during pregnancy: a meta-analysis. The Medical Journal of Australia, 208(1), 35–40.

Diamond, M. C., Johnson, R. E., & Ingham, C. (1971). Brain plasticity induced by environment and pregnancy. International Journal of Neuroscience, 2(4-5), 171–178.

Diamond, M. C., Krech, D., & Rosenzweig, M. R. (1964). The effects of an enriched environment on the histology of the rat cerebral cortex. Journal of Comparative Neurology, 123(1), 111–119.

Etkin, A., Egner, T., & Kalisch, R. (2011). Emotional processing in anterior cingulate and medial prefrontal cortex. Trends in cognitive sciences, 15(2), 85–93.

Everette, A., Fleming, D., Higgins, T., Tu, K., Bardi, M., Kinsley, C., & Lambert, K. (2007). Paternal experience enhances behavioral and neurobiological responsivity associated with affiliative and nurturing responses. International Behavioral Neuroscience Society, Rio de Janeiro.

Everette, A., Tu, K., Contino, R., Rima, B., Major, J., Conway, A., … Lambert, K. (2006). Plasticity of paternal responsiveness in two peromyscus species. International Behavioral Neuroscience Society, Whistler, BC.

Feldman, R. (2015). The adaptive human parental brain: implications for children’s social development. Trends Neurosci, 38(6), 387–399. doi:10.1016/j.tins.2015.04.004

Fleming, A. S., & Korsmit, M. (1996). Plasticity in the maternal circuit: effects of maternal experience on Fos-Lir in hypothalamic, limbic, and cortical structures in the postpartum rat. Behavioral Neuroscience, 110(3), 567.

Gatewood, J. D., Morgan, M. D., Eaton, M., McNamara, I. M., Stevens, L. F., Macbeth, A. H., … Kinsley, C. H. (2005). Motherhood mitigates aging-related decrements in learning and memory and positively affects brain aging in the rat. Brain Res Bull, 66(2), 91–98. doi:10.1016/j.brainresbull.2005.03.016

Henry, N. (2016). RN maternal newborn nursing: review module. Stilwell, KS: Assessment Technologies Institute. ISBN 9781565335691.

Hillerer, K. M., Neumann, I. D., Couillard-Despres, S., Aigner, L., & Slattery, D. A. (2014). Lactation-induced reduction in hippocampal neurogenesis is reversed by repeated stress exposure. Hippocampus, 24(6), 673–683. doi:10.1002/hipo.22258

Hoekzema, E., Barba-Muller, E., Pozzobon, C., Picado, M., Lucco, F., Garcia-Garcia, D., … Vilarroya, O. (2017). Pregnancy leads to long-lasting changes in human brain structure. Nature Neuroscience, 20(2), 287–296. doi:10.1038/nn.4458

Kim, P., Dufford, A. J., & Tribble, R. C. (2018). Cortical thickness variation of the maternal brain in the first 6 months postpartum: associations with parental self-efficacy. Brain Struct Funct, 223(7), 3267–3277. doi:10.1007/s00429-018-1688-z

Kim, P., Leckman, J. F., Mayes, L. C., Feldman, R., Wang, X., & Swain, J. E. (2010). The plasticity of human maternal brain: longitudinal changes in brain anatomy during the early postpartum period. Behavioural Neuroscience, 124(5), 695–700. doi:10.1037/a0020884

Kim, P., Rigo, P., Mayes, L. C., Feldman, R., Leckman, J. F., & Swain, J. E. (2014). Neural plasticity in fathers of human infants. Soc Neurosci, 9(5), 522–535. doi:10.1080/17470919.2014.933713

Kim, P., Strathearn, L., & Swain, J. E. (2016). The maternal brain and its plasticity in humans. Horm Behav, 77, 113–123. doi:10.1016/j.yhbeh.2015.08.001

Kinsley, C. H., & Amory-Meyer, E. (2011). Why the maternal brain? J Neuroendocrinol, 23(11), 974–983. doi:10.1111/j.1365-2826.2011.02194.x

Kinsley, C. H., Franssen, R. A., & Meyer, E. A. (2011). Reproductive experience may positively adjust the trajectory of senescence. In Behavioral Neurobiology of Aging (pp. 317–345): Springer.

Kinsley, C. H., & Lambert, K. G. (2008). Reproduction-induced neuroplasticity: natural behavioural and neuronal alterations associated with the production and care of offspring. J Neuroendocrinol, 20(4), 515–525. doi:10.1111/j.1365-2826.2008.01667.x

Kinsley, C. H., Madonia, L., Gifford, G. W., Tureski, K., Griffin, G. R., Lowry, C., … Lambert, K. G. (1999). Motherhood improves learning and memory. Nature, 402(6758), 137–138.

Kozorovitskiy, Y., Hughes, M., Lee, K., & Gould, E. (2006). Fatherhood affects dendritic spines and vasopressin V1a receptors in the primate prefrontal cortex. Nature Neuroscience, 9(9), 1094.

Laurent, H. K., & Ablow, J. C. (2011). A cry in the dark: depressed mothers show reduced neural activation to their own infant’s cry. Social Cognitive and Affective Neuroscience, 7(2), 125–134.

Lenzi, D., Trentini, C., Pantano, P., Macaluso, E., Iacoboni, M., Lenzi, G. L., & Ammaniti, M. (2009). Neural Basis of Maternal Communication and Emotional Expression Processing during Infant Preverbal Stage. Cerebral Cortex, 19(5), 1124–1133. doi:10.1093/cercor/bhn153

Macbeth, A. H., & Luine, V. N. (2010). Changes in anxiety and cognition due to reproductive experience: A review of data from rodent and human mothers. Neuroscience & Biobehavioral Reviews, 34(3), 452–467. doi:10.1016/j.neubiorev.2009.08.011

Maupin, A. N., Roginiel, A. C., Rutherford, H. J., & Mayes, L. C. (2016). A preliminary review of whether prior reproductive experience influences caregiving. New directions for child and adolescent development, 2016(153), 73–86.

McNeil, J. J., Woods, R. L., Nelson, M. R., Murray, A. M., Reid, C. M., Kirpach, B., … Tonkin, A. M. (2017). Baseline characteristics of participants in the ASPREE (Aspirin in Reducing Events in the Elderly) study. Journals of Gerontology Series A: Biomedical Sciences and Medical Sciences, 72(11), 1586–1593.

Musser, E. D., Kaiser-Laurent, H., & Ablow, J. C. (2012). The neural correlates of maternal sensitivity: an fMRI study. Developmental cognitive neuroscience, 2(4), 428–436.

Nishitani, S., Kuwamoto, S., Takahira, A., Miyamura, T., & Shinohara, K. (2014). Maternal prefrontal cortex activation by newborn infant odors. Chemical senses, 39(3), 195& 202.

Numan, M., Corodimas, K. P., Numan, M. J., Factor, E. M., & Piers, W. D. (1988). Axon-sparing lesions of the preoptic region and substantia innominata disrupt maternal behavior in rats. Behavioral Neuroscience, 102(3), 381.

Numan, M., & Insel, T. R. (2003). Motivational models of the onset and maintenance of maternal behavior and maternal aggression. The neurobiology of parental behavior, 69–106.

Numan, M., & Numan, M. J. (1991). Preoptic-brainstem connections and maternal behavior in rats. Behavioral Neuroscience, 105(6), 1013.

Oatridge, A., Holdcroft, A., Saeed, N., Hajnal, J. V., Puri, B. K., Fusi, L., & Bydder, G. M. (2002). Change in brain size during and after pregnancy: study in healthy women and women with preeclampsia. American Journal of Neuroradiology, 23(1), 19–26.

Olson, I. R., Plotzker, A., & Ezzyat, Y. (2007). The enigmatic temporal pole: a review of findings on social and emotional processing. Brain, 130(7), 1718–1731.

Radloff, L. S. (1977). The CES-D scale: A self-report depression scale for research in the general population. Applied psychological measurement, 1(3), 385–401.

Rosenblatt, J. S. (1994). Psychobiology of maternal behavior: contribution to the clinical understanding of maternal behavior among humans. Acta paediatrica, 83(397), 3–8.

Ross, T. P. (2003). The reliability of cluster and switch scores for the Controlled Oral Word Association Test. Archives of Clinical Neuropsychology, 18(2), 153–164.

Seo, S. W., Im, K., Lee, J.-M., Kim, S. T., Ahn, H. J., Go, S. M., … Na, D. L. (2011). Effects of demographic factors on cortical thickness in Alzheimer’s disease. Neurobiology of aging, 32(2), 200–209.

Smith, A. (1982). Symbol digit modalities test: Western Psychological Services Los Angeles, CA.

Speisman, R. B., Kumar, A., Rani, A., Pastoriza, J. M., Severance, J. E., Foster, T. C., & Ormerod, B. K. (2013). Environmental enrichment restores neurogenesis and rapid acquisition in aged rats. Neurobiol Aging, 34(1), 263–274. doi:10.1016/j.neurobiolaging.2012.05.023

Stolzenberg, D. S., & Champagne, F. A. (2016). Hormonal and non-hormonal bases of maternal behavior: The role of experience and epigenetic mechanisms. Horm Behav, 77, 204–210. doi:10.1016/j.yhbeh.2015.07.005

Storey, A. E., Walsh, C. J., Quinton, R. L., & Wynne-Edwards, K. E. (2000). Hormonal correlates of paternal responsiveness in new and expectant fathers. Evolution and Human Behavior, 21(2), 79–95.

Swain, J. E., Lorberbaum, J. P., Kose, S., & Strathearn, L. (2007). Brain basis of early parent-infant interactions: psychology, physiology, and in vivo functional neuroimaging studies. Journal of Child Psychology and Psychiatry, 48(3-4), 262–287. doi:10.1111/j.1469-7610.2007.01731.x

Swain, J. E., Tasgin, E., Mayes, L. C., Feldman, R., Constable, R. T., & Leckman, J. F. (2008). Maternal brain response to own baby-cry is affected by cesarean section delivery. J Child Psychol Psychiatry, 49(10), 1042–1052. doi:10.1111/j.1469- 7610.2008.01963.x

Teng, E., & Chui, H. (1987). The modified mini-mental state examination (3MS). Can J Psychiatry, 41(2), 114–121.

Troyer, A. K., Leach, L., & Strauss, E. (2006). Aging and response inhibition: Normative data for the Victoria Stroop Test. Aging, Neuropsychology, and Cognition, 13(1), 20–35.

Veit, R., Kullmann, S., Heni, M., Machann, J., Häring, H.-U., Fritsche, A., & Preissl, H. (2014). Reduced cortical thickness associated with visceral fat and BMI. NeuroImage: Clinical, 6, 307–311.

Volkmar, F. R., & Greenough, W. T. (1972). Rearing complexity affects branching of dendrites in the visual cortex of the rat. Science, 176(4042), 1445–1447.

Wager, T. D., Davidson, M. L., Hughes, B. L., Lindquist, M. A., & Ochsner, K. N. (2008). Prefrontal-subcortical pathways mediating successful emotion regulation. Neuron, 59(6), 1037–1050.

Ward, S. A., Raniga, P., Ferris, N. J., Woods, R. L., Storey, E., Bailey, M. J., … on behalf of the ASPREE Investigator Group. (2017). ASPREE-NEURO study protocol: A randomized controlled trial to determine the effect of low-dose aspirin on cerebral microbleeds, white matter hyperintensities, cognition, and stroke in the healthy elderly. Int J Stroke, 12(1), 108–113. doi:10.1177/1747493016669848

